# Genome-wide diversity in lowland and highland maize landraces from southern South America: population genetics insights to assist conservation

**DOI:** 10.1101/2024.02.02.578655

**Authors:** Pia Guadalupe Dominguez, Angela Veronica Gutierrez, Monica Irina Fass, Carla Valeria Filippi, Pablo Vera, Andrea Puebla, Raquel Alicia Defacio, Norma Beatriz Paniego, Veronica Viviana Lia

**Author notes:** These authors contributed equally to this work (Pia Guadalupe Dominguez, Angela Veronica Gutierrez). **E-mails:** PGD, AVG, MIF, CVF, PV, AP, RAD, NBP, VVL.

## Abstract

Maize (*Zea mays* ssp. *mays L.*) landraces are traditional American crops with high genetic variability that conform a source of original alleles for conventional maize breeding. Northern Argentina, one the southernmost regions of traditional maize cultivation in the Americas, harbours around 57 races traditionally grown in two regions with contrasting environmental conditions, namely the Andean mountains in the Northwest and the tropical grasslands and Atlantic Forest in the Northeast. These races encounter diverse threats to their genetic diversity and persistence in their regions of origin, with climate change standing out as one of the major challenges. In this work, we use genome-wide SNPs derived from ddRADseq to study the genetic diversity of individuals representing the five groups previously described for this area. This allowed us to distinguish two clearly differentiated gene pools, the Highland Northwestern maize (HNWA) and the Floury Northeastern maize (FNEA). Subsequently, we employed Essential Biodiversity Variables at the genetic level, as proposed by the Group on Earth Observations Biodiversity Observation Network (GEO BON), to evaluate the conservation status of these two groups. This assessment encompassed genetic diversity (Pi), inbreeding coefficient (F), and effective population size (Ne). FNEA showed low Ne values and high F values, while HNWA showed low Ne values and low Pi values, indicating that further genetic erosion is imminent for these landraces. Outlier detection methods allowed identification of putative adaptive genomic regions, consistent with previously reported flowering-time loci and chromosomal regions displaying introgression from the teosinte *Zea mays* ssp. *mexicana*. Finally, species distribution models were obtained for two future climate scenarios, showing a notable reduction in the potential planting area of HNWA and a shift in the cultivation areas of FNEA. Taken together, these results suggest that maize landraces from Northern Argentina may not be able to cope with climate change. Therefore, active conservation policies are advisable.

## Introduction

Maize landraces are varieties that have been grown by local communities throughout the Americas since pre-Columbian times (Gupta et al., 2020). They differ from commercial hybrids in that they are open-pollinated, and cultivated through traditional methods (Mercer et al., 2008; Casañas et al., 2017; Gupta et al., 2020). The cultivation characteristics of these varieties include cross-pollination between fields, seed exchange by farmers, and selection by both agricultural management and environmental conditions (Mercer et al., 2008; Casañas et al., 2017). Due to this, landraces often have high genetic variability and constitute a valuable source of original alleles for breeding. On the other hand, commercial hybrids capture only a small fraction of this variation, because of the use of a limited set of landraces in breeding programs (Hufford et al., 2012; Smith et al., 2017). Moreover, the replacement of landraces with more productive, but genetically uniform, commercial germplasm has led to significant genetic erosion (Dwivedi et al., 2016, Heck et al., 2020, Gupta et al., 2020). Therefore, active landrace conservation actions are essential to preserve the genetic and phenotypic variability of this crop.

The Group on Earth Observations Biodiversity Observation Network (GEO BON; https://geobon.org/) has defined the Essential Biodiversity Variables (EBVs) as a set of variables of different origin that serve to capture critical scales and dimensions related to biodiversity, including how biodiversity is geographically distributed and how it varies over time (Pereira et al., 2013; Brummitt et al., 2017; Navarro et al., 2017; Schmeller et al., 2017; Hoban et al., 2022). At the genetic level, Hoban et al. (2022) proposed to evaluate four EBVs: genetic diversity, genetic differentiation, inbreeding and effective population size, which provide information on genetic variation at different levels (within populations, between populations, within individuals, and change in genetic diversity due to drift, respectively) using a single genomic data set. EBVs encompass metrics that can be used to forecast the status and trends of genetic diversity, which is the cornerstone of species resilience, and essential to their ability to adapt to environmental conditions (Hoban et al., 2022). Although the concept of EBV is usually applied to natural populations or invasive species (Hoban et al., 2022), these metrics could also be applied to domesticated species such as maize given that EBVs respond to both natural and anthropogenic drivers.

Climate change is currently one of the main threats to crop species diversity, making germplasm conservation one of the most pressing present-day challenges (Gupta et al., 2020). Commercial maize production is estimated to fall by 50% with a 4°C temperature increase and by 10% with a 2°C temperature increase in major maize producing countries (Tigchelaar et al., 2018). Landraces are characterised by being locally adapted, i.e., by presenting greater fitness in their native habitats than in other environments (Savolainen et al., 2013). Under a climate change scenario, the only possibilities for landraces to survive in their original locations are either by evolving via selection upon standing variation or through plasticity (Cang et al., 2016). However, Cang et al. (2016) estimated that the speed of climate change is 5,000 times faster than the adaptive capacity of 230 species of the *Gramineae* family. This suggests that rapid adaptation to changing conditions in local environments is not likely to happen, implying that climate change may significantly affect maize landraces too. Understanding how local germplasm has adapted to its surroundings can help lessen the potential of diversity reductions. Thus, in addition to EBVs, focusing on adaptive variation adds a significant aspect to conservation considerations since identifying genes under selection may help quantify the extent of local adaptation and provide information on the molecular processes behind phenotypic divergence.

Northern Argentina is one of the southernmost regions of maize landrace cultivation in South America and it has been proposed as an ancient contact zone between Andean and Tropical lowland germplasm (Vigoroux et al., 2008; Tenaillon and Charcosset, 2011). This area harbours ca. 57 maize landraces and encompasses two clearly differentiated agroecosystems: the Northwest, and the Northeast (Bracco et al., 2012; Melchiorre et al., 2017; Realini et al., 2018). In Northwestern Argentina (NWA), maize cultivation extends to an altitude of ca. 4,000 meters above sea level (m.a.s.l.), daily temperature ranges are large, precipitations are below 350 mm/year, oxygen pressure is low, soil nutrients are scarce and radiation indices are high (Rivas et al., 2022). By contrast, altitude in Northeastern Argentina (NEA) does not exceed 800 m.a.s.l. while climate is subtropical, with average annual temperature between 15 and 23 °C and annual precipitation between 1,000 and 2,000 mm. Soils in NEA are clayish with limiting components (nitrogen, phosphorus, organic matter), low pH and low to medium fertility (Heck et al., 2020).

Bracco et al. (2016) found significant molecular differentiation between NWA and NEA landraces, and identified three genetic groups: NWA maize, Floury Northeastern maize (FNEA) and Northeastern Popcorns (PNEA). More recently, microsatellite analysis of NWA landraces revealed that there is an altitude-associated genetic structure, with two main genetic pools: Highland Northwestern maize or HNWA, cultivated at more than 2,000 m.a.s.l., and Lowland Northwestern maize or LNWA, cultivated below 2,000 m.a.s.l. (Rivas et al., 2022). Additionally, a third NWA group, the Northwestern Popcorns (PNWA), was recognized by Lia et al. (2009). Previous studies showed that HNWA are associated with Andean landraces and that FNEA represents a unique, locally adapted gene pool, with no clear connections to any other lowland maize from South America (Lia et al., 2009; Bracco et al., 2016; Lopez et al., 2021). Similarly, the origins and affiliations of LNWA remain unknown. Overall, the complex structuring of genetic diversity suggests that further efforts are still needed to delineate significant units and effectively assist conservation.

In this work, we use genome-wide molecular markers derived from ddRADseq and the genetic EBVs proposed by Hoban et al. (2022) to assess the conservation status of maize landraces from NWA and NEA. In addition, to test for evidence of adaptive divergence we searched for selection signals. Finally, we used two future climate scenarios to perform Bayesian modelling of species distribution. The results of this work suggest that the long-term diversity of maize landraces of Northern Argentina is compromised, and that more active conservation policies are advisable.

## Materials and methods

### Plant Material

A set of 87 maize individuals representative of the genetic and morphological groups previously identified for the Northeast and Northwest of Argentina were obtained from the “Banco Activo de Germoplasma INTA Pergamino” (BAP; INTA, Pergamino, Buenos Aires, Argentina) and from the “N.I. Vavilov” Plant Genetic Resource Laboratory, Faculty of Agronomy, University of Buenos Aires. General characteristics of the accessions, including ID, racial classification, and collection site, are given in Figure 1A and Supplementary Table 1. A priori group assignment is based on the analysis of microsatellite data according to Lia et al. (2009), Bracco et al. (2016), López et al. (2021), and Rivas et al. (2022) (Supplementary Table 1). The map was made with QGIS v3.16.16-Hannover (https://qgis.org/en/site/), employing a 1:50m political map from Natural Earth (https://www.naturalearthdata.com/) and a 5-minute latitude/longitude grid digital elevation model from the European Environment Agency (https://data.europa.eu/data/datasets/data_world-digital-elevation-model-etopo5?locale=es).

**Figure 1.**
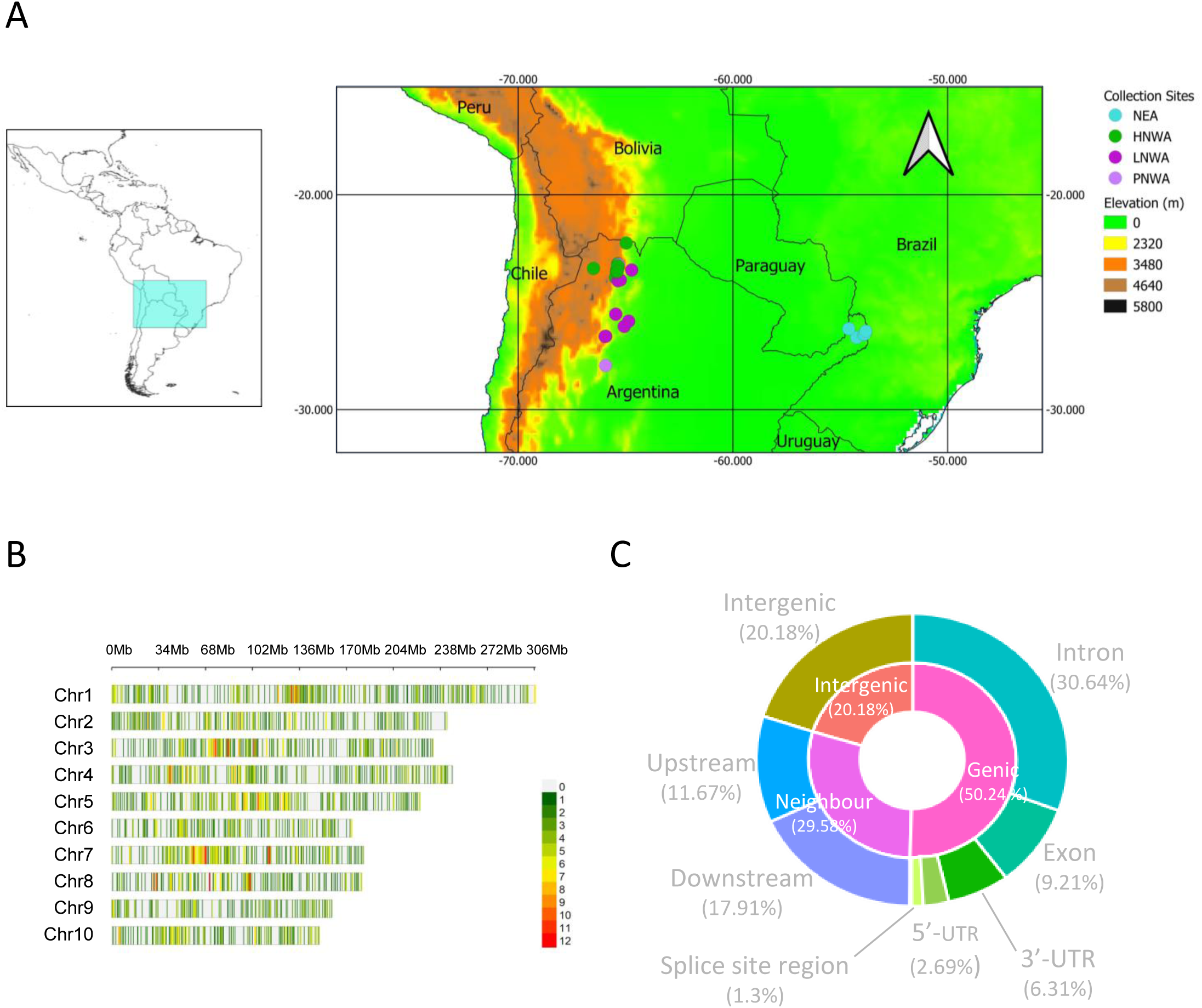
Characterisation of maize landrace accessions from Northern Argentina by ddRADseq. A) Collection sites of the individuals included in this study. The map was made with QGIS. The list of individuals is in Supplementary table 1. B) Distribution of the SNPs detected in the chromosomes. The plot was made with CMplot (Yin et al. 2021). The colours indicate the number of SNPs in a 1 Mbp window. C) Summary of the annotation of the SNP matrix according to the region performed with SnpEff (Cingolani et al. 2011). HNWA: Highland maize of Northwestern Argentina. LNWA: Lowland maize of Western Argentina. PNWA: Popcorn of Northwestern Argentina. NEA: Northeastern Argentina (Floury maize of Northeastern Argentina and Popcorn of Northeastern Argentina). Total number of individuals: 87.

### DNA Extraction

Plants were germinated in a greenhouse under controlled conditions (80% relative humidity; 200 mmol PAR s-1m-2; 16 h of light/8 h of darkness). DNA was extracted with the protocol by Dellaporta et al. (1983) from 100 mg of fresh leaves. The quality of the DNA was checked using a NanoDrop1000 (DNA quality criterion by absorbance: A260/A280 > 1.8 and A260/A230 ≈ 1.8-2.2) and through runs on 0.8% agarose gels. DNA was quantified using a Qubit 2.0 fluorometer (Thermo Fisher Scientific).

### DNA Sequencing

The preparation of the genomic libraries was carried out using the protocol developed by Aguirre et al. (2019). Briefly, DNA samples were digested with two digestion enzymes, one of rare cleavage and one of frequent cleavage (SphI + MboI). Adapters (4-9 bp) published by Peterson et al. (2014) were ligated to the digested fragments. The reactions were incubated for one hour at 23 °C, followed by an additional one-hour incubation at 20 °C. Ligations from all samples were mixed in equal DNA amounts in pools of 22-24 individuals, concentrated, and finally purified with 1X Ampure XP beads per group. Next, an automated size selection (one for each pool) was performed on a 2% agarose cassette in SAGE ELF (Sage Science, Inc., Beverly, MA, USA). 450-bp fragments were retained and subsequently purified with 0.8X AMPure beads (Beckman Coulter, Indianapolis, USA). Finally, PCRs were carried out for each of the pools employing dual-indexed primers (Lange et al., 2014). The four pools were put together. Low-depth sequencing was performed on a MySeq Illumina (Albany, USA) equipment in the Genomics Unit of the IABIMO (Hurlingham, Buenos Aires, Argentina) to verify the correct assembly of the library. The samples were sent to the International Maize and Wheat Improvement Center (CYMMIT, El Batán, Texcoco, Mexico), where they were sequenced on an Illumina Novaseq (Albany, USA) device with paired-end readings (2×150 bp).

### ddRADseq bioinformatics analysis

Raw reads were curated for quality in Stacks v1.42 (Catchen et al., 2013). Barcodes were removed and reads were trimmed to 150 bp. SNP calling was also performed with Stacks v1.42. The parameters used were: -m 3 (minimum depth of coverage), - M 2 (distance allowed between stacks), -n 3 (distance allowed between catalog loci). Reads of each sample were mapped against the V4 version of maize B73 reference genome (https://www.maizegdb.org/genome/assembly/Zm-B73-REFERENCE-GRAMENE-4.0) with Bowtie 2 (Langmead et al., 2012). The resulting vcf file was filtered with VCFtools (Danecek et al., 2011). Only sites fulfilling the following requirements were retained: a maximum proportion of missing data of 35% (--max-missing 0.65); a minimum number of times that an allele appears over all individuals at a given site equal to 4 (--mac 4); a mean depth value greater than or equal to 8 per individual (--minDP 8); a minimum distance between sites equal to 200 bp (--thin200). The unfiltered and filtered vcf files are provided in Supplementary tables 2 and 3, respectively. The imputation of the filtered vcf file was carried out with Beagle (Browning et al., 2018). The filtered imputed vcf file is found in Supplementary table 4. The genomic variant annotation was performed with SnpEff (Cingolani et al., 2012). The filtered, annotated, and imputed vcf file is found in Supplementary table 5. This vcf file was used for all subsequent analyses. The graph of the SNP density was plotted with the CMplot package (Yin et al., 2021) in R (https://www.r-project.org/).

### Population structure analyses

A neighbor joining (NJ) phylogenetic tree based on Euclidean distances was built with Tassel (Bradbury et al., 2007) and graphed with Itol (https://itol.embl.de/). The principal component analysis (PCA) was performed with the Adegenet package version 2.1.10 in R (Jombart et al., 2008). The discriminant analysis of principal components (DAPC) was performed using the Adegenet package in R (Jombart et al., 2008). The “find.clusters” function was used to find the optimal number of clusters (k) to describe the data employing the BIC values criteria. The DAPC itself was implemented with the “xvalDapc” function using the previously inferred k groups and cross-validation to define the number of PCs. A Bayesian analysis of population structure was performed with the STRUCTURE software employing the admixture model with correlated allele frequencies (Pritchard et al., 2000). Between 2 and 6 clusters (Ks) were evaluated running 3 times each K (burn-in: 50,000; iterations: 100,000). The deltaK method (Evanno et al., 2005) was used to determine the most probable K through the Structure Harvester program (Earl et al., 2012). The allele frequency divergence estimate given by the software was used to measure the differentiation between STRUCTURE groups.

### Characterisation of potential conservation units

Based on the findings of the various population structure analyses, two groups, HNWA and FNEA, were chosen for further investigation using EBVs, genome scans of selection, and habitat distribution modelling. Only those individuals that were unequivocally assigned to each genetic cluster by the STRUCTURE and DAPC methods were considered for further analyses (membership coefficients or assignment probabilities > 0.75, respectively) (Supplementary Table 1).

### Linkage disequilibrium

Linkage disequilibrium was calculated as the squared allele frequency correlation (r^2^) employing the --geno-r2 option of VCFtools (Danecek et al., 2011). The expected decay of linkage disequilibrium (r^2^) with physical distance was modelled for each chromosome using Hill and Weir’s equation (1988) on the basis of the script developed by Marroni et al. (2011).

### Estimation of EBVs

Nucleotide diversity (Pi) per site, inbreeding coefficient (F), Hardy-Weinberg equilibrium and fixation index per site (Fst, Weir and Cockerham, 1984) were computed with VCFtools (Danecek et al., 2011) using the --site-pi, --het, --hardy and --weir-fst-pop functions, respectively. The nucleotide diversity per site as calculated with VCFtools is equivalent to the expected heterozygosity. Graphs were plotted with ggplot2 in R (Wickham, 2016), with the Pi per site graphs being Loess smoothing plotted. The Hardy-Weinberg equilibrium plots were made with the CMplot package in R (Yin et al., 2021). The effective population size was estimated employing the linkage disequilibrium method implemented in NeEstimator v.2.0 (Do et al., 2014). In accordance with the suggestions of Hoban et al. (2022), we employed Pi per site as a proxy for genetic diversity, Fst as a measure of genetic differentiation, F to evaluate individual inbreeding and the LD estimate of Ne to assess the contemporary effective population size.

### Analysis of outlier loci

The genomic signatures of selection were searched for with BayPass version 2.4 (Gautier, 2015), which accounts for the shared ancestry and population structure within the dataset by generating a covariance matrix of allele frequencies (Ω). SNPs under selection were detected employing the core model with default options. The identification of outliers was based on a calibration procedure of XtX values using pseudo-observed datasets (PODs) of 3,500 SNPs and a 1% threshold. XtX values are analogous to Fst but formally corrected by the covariance matrix. The Manhattan plot showing the XtX values against SNP chromosomal positions was generated with the CMplot package in R (Yin et al., 2021). Genes 1 Mb upstream or downstream of SNPs under positive selection were considered as candidates to be associated with them. The gff3 file of the V4 version of the maize B73 reference genome (https://www.maizegdb.org/genome/assembly/Zm-B73-REFERENCE-GRAMENE-4.0) was filtered by the chromosome in which the SNP was found using an awk command. Filters included the interval of 1 Mb up- and downstream of the position of the outlier SNP, and the “gene” category of each feature. If available, the annotation of each gene was considered. Otherwise, the annotation of those genes containing outlier SNPs was inferred by similarity to genes from other species.

### Habitat suitability modelling

The geographical distribution of the FNEA and HWNA groups was modelled with MaxEnt version 3.4.4 (Phillips et al., 2006) employing historical bioclimate variables (period 1970-2000) and elevation data. Briefly, a total of 158 geographically unique records were used, 25 for FNEA and 133 for HWNA. Occurrence records include geographical coordinates of the individuals used in this study and those reported for other individuals from the same genetic groups by Bracco et al. (2016) (Supplementary table 6). Models were generated using 20,000 background points from all over the world, using hinge features only and default regularisation parameters as recommended by Bracco et al. (2016). Model performance was assessed using the area under the receiver operating characteristic curve (AUC) for both training and testing data sets. To account for the differences in sample sizes, ten and four-fold cross-validation were employed for HNWA and FNEA, respectively, to estimate errors around fitted functions and predictive performance on held-out data. The contribution of each variable to model improvement throughout the training process (percentage of contribution) and jackknife tests implemented in MaxEnt were used to determine variable relevance. The models were subsequently projected to two future climate scenarios, CNRM-CM6-1 (Voldoire et al., 2019) and MRI-ESM2-0 (Yukimoto et al., 2019), for the period 2081-2100 and under four CO_2_ emission scenarios (SSP5-8.5, SSP3-8.7, SSP2-4.5, SSP1-2.6). These two future climate scenario models were chosen because they are in the middle zone of the high sensitivity models (CNRM-CM6-1) and in the middle zone of the standard sensitivity models (MRI-ESM2-0) of WorldClimb. All bioclimate variables and elevation data have a 2.5-minute spatial resolution and were retrieved from WorldClim (https://www.worldclim.org/data/cmip6/cmip6climate.html). Pairwise comparisons of model predictions were carried out by calculating the Schoener’s D (Schoener, 1968) and the I statistic (Warren et al., 2008) in ENMtools 1.3 (Warren et al., 2021).

## Results

### SNP discovery and annotation

Eighty-seven individuals representative of the five genetic and morphological groups previously identified for northern Argentina (i.e., HNWA: Highland maize from Northwestern Argentina; LNWA: Lowland maize from Western Argentina; PNWA: Popcorn from Northwestern Argentina; FNEA: Floury maize from Northeastern Argentina; and PNEA: Popcorn from Northeastern Argentina) were sequenced through ddRADseq (Figure 1A). A total of 3,529 SNPs distributed along the 10 maize chromosomes were obtained after filtering and imputation of the raw data matrix (Figure 1B). Functional annotation of the SNPs indicated that only a small proportion of the variants was found within exons (9.21%), with the highest percentages predicted as intronic (30.64%), intergenic (20.18%), or located downstream of genes (17.91%) (Figure 1C).

### Analysis of population structure

Both the Neighbor-Joining tree and the PCA show two clear groups, one formed mainly by HNWA individuals and the other by FNEA individuals (Figure 2A and B). PNWA and PNEA individuals tend to cluster together within each group but closely with LNWA individuals, which occupy an intermediate position in both the network and PCA biplot. Therefore, the distinction of these three groups (PNEA, LNWA, PNWA) is less clear. It is noteworthy that, among the five LNWA individuals demonstrating a close affinity to the FNEA group, three were morphologically classified as Avati morotí (Supplementary table 1), a race indigenous to the NEA region.

**Figure 2.**
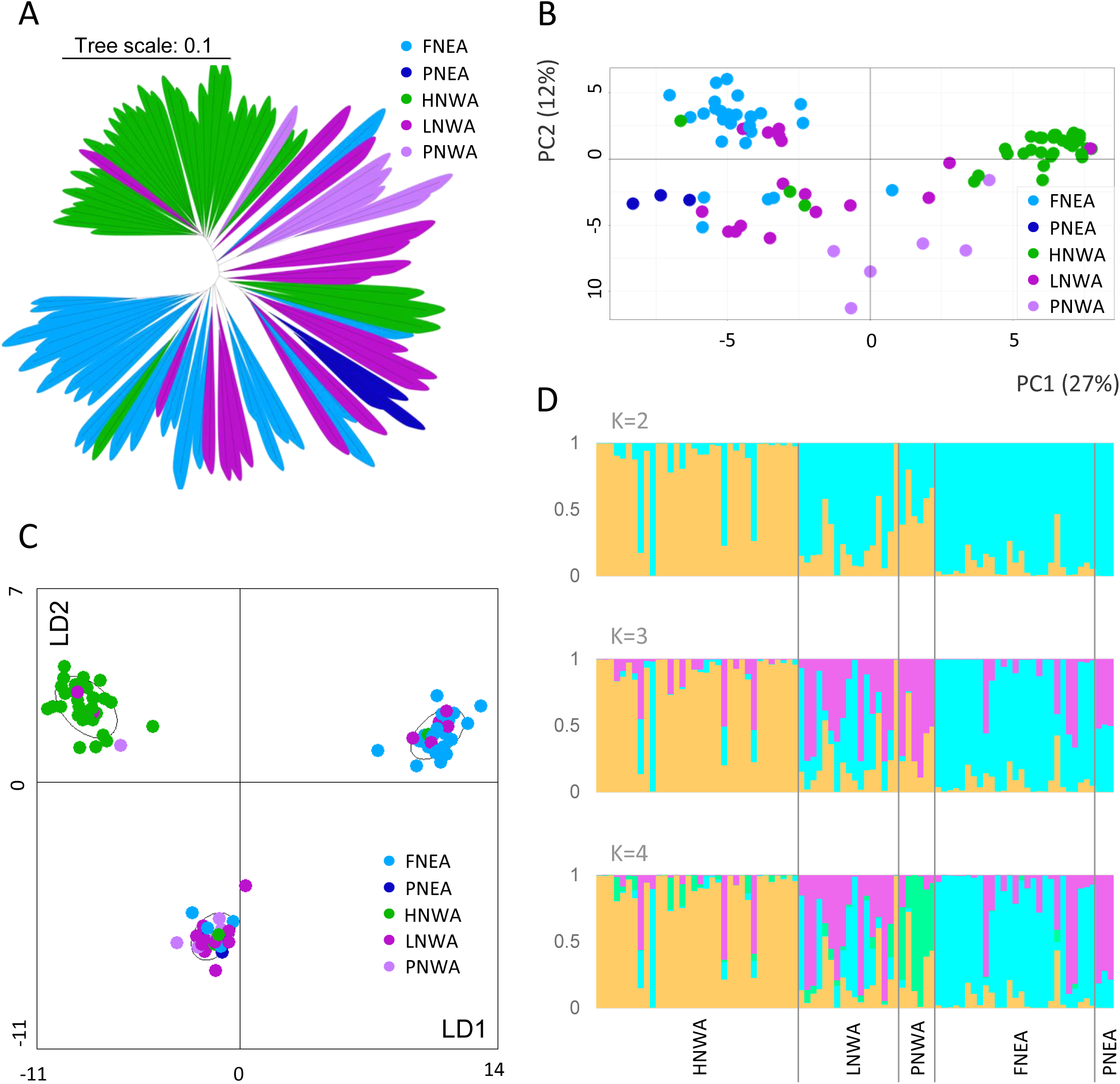
Analysis of the population structure. A) Neighbor Joining tree employing Euclidean distance (Bradbury et al. 2007). B) Principal component analysis (PCA) performed with Adegenet in R (Jombart et al. 2008). PC: Principal component. C) Discriminant Analysis of Principal Components performed with Adegenet in R, k=3. LD: Linear Discriminant Axis. D) Bayesian analysis performed with STRUCTURE (Pritchard et al. 2000), K=2-4. Individuals were classified *a priori* according to Lia *et al*. (2009), Bracco *et al*. (2016), López *et al*. (2021), and Rivas *et al*. (2022) : HNWA (Highland maize of Northwestern Argentina), LNWA (Lowland maize of Western Argentina), FNEA (Floury maize of Northeastern Argentina), PNEA (Popcorn of Northeastern Argentina), and PNWA (Popcorn of Northwestern Argentina).

Based on the BIC criterion, the k-means algorithm identified k=2 and k=3 as the two most probable numbers of groups for the DAPC (Figure 2C, Supplementary Figure 1). At k=3, one cluster was enriched with FNEA, another with HNWA, and a third with individuals from every a priori group (Figure 2C, Supplementary Figure 1D), whereas at k=2, the discriminant function mostly distinguished HNWA from the remaining individuals (Supplementary Figure 1E and F).

In agreement with the DAPC, STRUCTURE analysis with K=2 (the most probable K according to the delta-K method) shows that one cluster is mainly made up of HNWA individuals (orange), while the second cluster is made up of the rest of the individuals (light blue), with PNWA receiving almost equal contributions from both clusters (Figure 2D). With K=3, one group consists of HNWA individuals (orange), another group consists of NEA (FNEA and PNEA) individuals (light blue), and the third group consists mainly of individuals from LNWA and PNWA (pink) (Figure 2D). When K=4, there are two groups formed mainly by HNWA individuals (orange) and FNEA individuals (light blue), respectively, while the pink group is formed mainly by relatively admixed individuals from LNWA and PNEA (Figure 2D). In turn, PNWA individuals separate into an independent group (green) (Figure 2D). Allele frequency divergence for the inferred clusters varied from 0.0263 (light blue vs. pink) to 0.072 (orange vs. green) (Supplementary Table 7, K=4). Ascending in magnitude, the genetic drift parameters for the pink, light blue, orange, and green clusters— representing their divergence from a common hypothetical ancestor—were 0.161, 0.207, 0.362 and 0.442, respectively.

Collectively, these findings show that HNWA and FNEA consistently emerge as the two predominant groups, implying the presence of at least two distinct conservation units in Northern Argentina. Due to the limited sample sizes of PNEA and PNWA, along with the apparent heterogeneity within LNWA, these groups were not considered in subsequent analyses.

### Linkage disequilibrium

Linkage disequilibrium decay was examined for each of the two main groups identified in the previous analyses (HNWA and FNEA) (Figure 3). Both average and single chromosome estimates showed a more rapid decay for HNWA than for FNEA, with r^2^ reaching 0.1 at approximately 2.2 and 2.9 MB, respectively (Figure 3). In line with this, average r^2^ values overall chromosomes were 0.046 for HNWA and 0.058 for FNEA.

**Figure 3.**
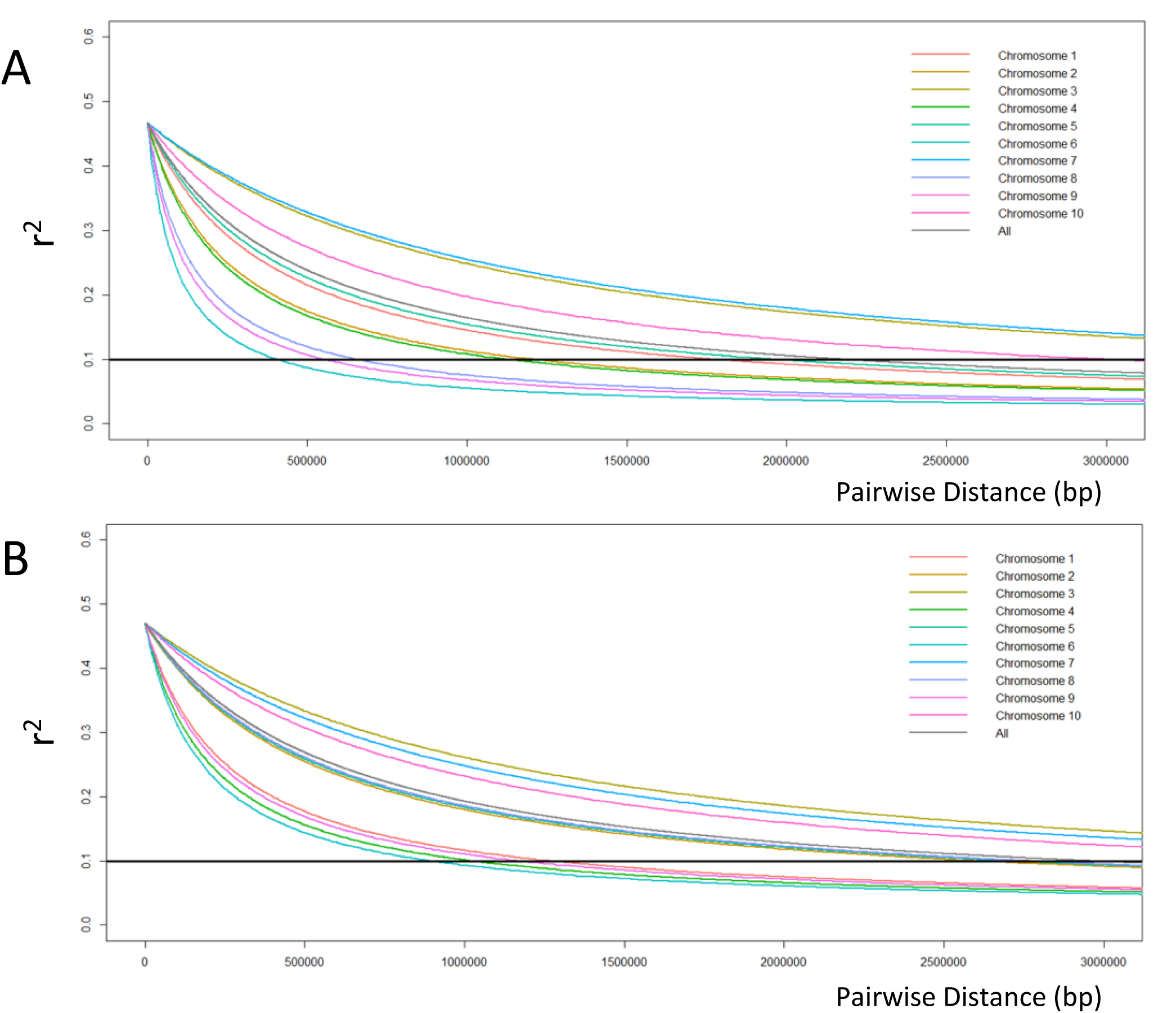
Decay of linkage disequilibrium calculated as the squared allele frequency correlation (r^2^). r^2^ is plotted against the physical distance between markers of (A) Highland maize of Northwestern Argentina, and (B) Floury maize of Northeastern Argentina. The cut-off line is plotted at r^2^ = 0.1. The fitting of the curves was done according to the Hill and Weir’s equation (1988).

### Nucleotide diversity (Pi), inbreeding coefficient (F) and effective population size (Ne)

Population diversity indices were estimated for the entire set of individuals (N=87), as well as for HNWA and FNEA. Patterns of variation along chromosomes were consistent across the three groups, however Pi values per site tended to be lower in HNWA (average Pi per site= 0.173, Figure 4A), indicating less genetic variability than in FNEA (average Pi per site=0.205). Tests of Hardy-Weinberg proportions revealed that only a few SNP loci deviated from panmixia in both the HNWA and FNEA groups, as expected for outcrossing species (Supplementary Figure 2). When the total number of individuals was considered, the proportion of loci with homozygote excess rose because of population sub-structuring. For its part, estimates of inbreeding coefficients based on individual heterozygosity (F_H_) showed that consanguinity tended to be higher in FNEA than in HWNA individuals, with distributions centred around F_H_=0.25 and F_H_=0.12, respectively (Figure 4B). Negative F_H_ values imply that the parents of those individuals were less related than expected under random mating, a phenomenon that may be frequently encountered in maize because of human-mediated introductions of exogenous germplasm. In terms of effective population size, the FNEA group exhibited contemporary Ne values of 51.3, 65.2, and 65.2 individuals, depending on the MAF (minimum allele frequency) thresholds of 0.05, 0.02, and 0.01, respectively (Figure 4C). Conversely, the HNWA group presented Ne values of 245.7, 181.1, and 143.9 for each MAF (Figure 4C).

**Figure 4.**
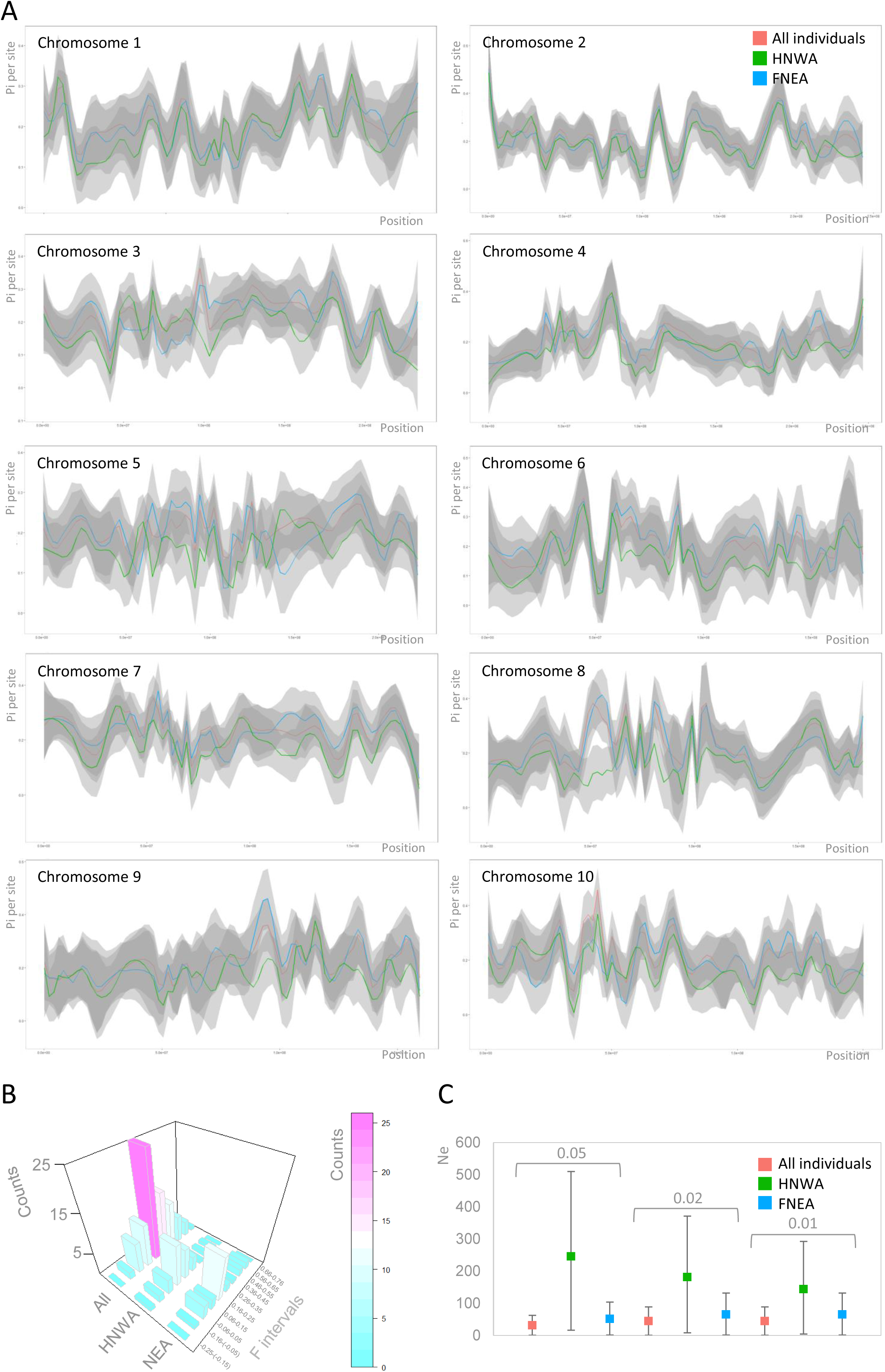
Diversity and effective population sizes. A) Nucleotide diversity per site (Pi or π) for each chromosome computed with VCFtools (Danecek et al., 2011). Values were adjusted by nonparametric local regression (LOESS). B) Histogram showing the inbreeding coefficient (F) calculated with VCFtools. C) Effective population size (Ne) estimated employing the linkage disequilibrium method implemented in NeEstimator v.2.0 (Do et al., 2014). Squares indicate the arithmetic mean, while the bars indicate 95% confidence intervals. Minimum allele frequency used: 0.05, 0.02 and 0.01. HNWA: Highland maize of Northwestern Argentina. FNEA: Floury maize of Northeastern Argentina.

### Genetic differentiation and outlier loci

Analysis of genetic differentiation between HNWA and FNEA revealed an average Fst value of 0.07. The distribution of Fst values across all chromosomes was generally uniform, although chromosomes 3, 7, and 10 displayed slightly larger interquartile ranges (Figure 5A). To delve deeper into the nature and distribution of adaptive variation, we conducted a search for outlier loci using the BayPass program, identifying 56 loci that exhibited signatures of directional selection and can be potentially associated with local adaptation (Figure 5B and Supplementary Table 8). Annotation of these SNPs revealed that the majority were located within intergenic regions (Supplementary Table 8A), though no enrichment was observed for outliers in this category compared to the complete data matrix (Fisher exact test, p > 0.05). Among the seven outlier SNPs located within gene bodies, we identified candidates associated with flowering time and stress responses (Supplementary Table 8A). In addition to the outlier SNPs identified within genes, three chromosome regions present a notable abundance of outlier SNPs. Seven of the 56 outlier SNPs, representing 5 ddRAD loci, were situated within a 1 MB region proximal to the centromere on chromosome 3, while two larger blocks were detected in chromosomes 7 and 10 (Supplementary Table 8B). Gene models and annotations within to 2 MB windows around outlier SNPs are provided in Supplementary Table 8. This window size was selected taking into consideration the observed extent of LD.

**Figure 5.**
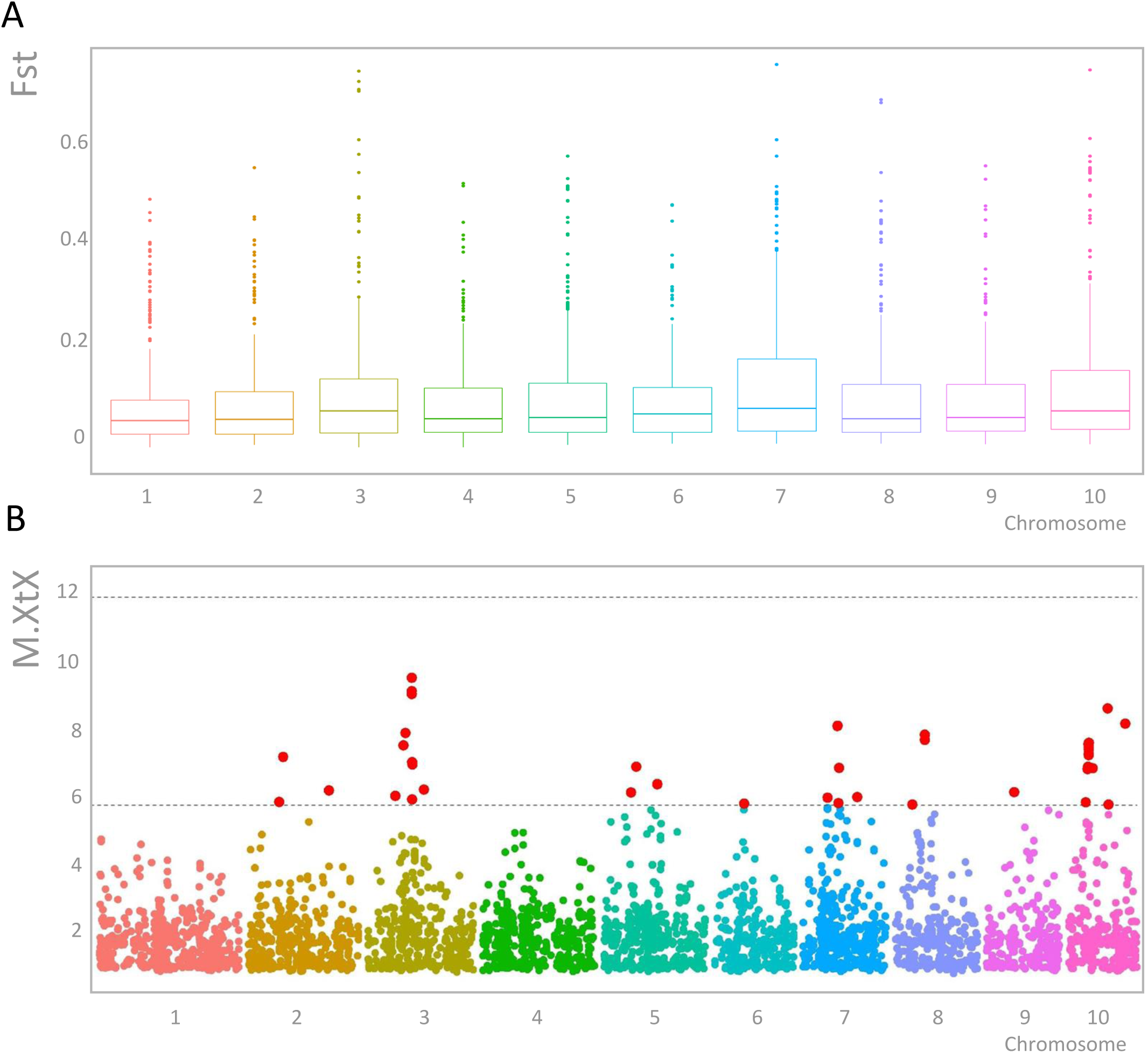
Detection of genomic signatures of selection between Floury maize of Northeastern Argentina (FNEA) and Highland maize of Northwestern Argentina (HNWA). A) Box-plot of Fst values per chromosome obtained with VCFtools (Danecek et al. 2011). B) Determination of outlier loci with BayPass (Gautier, 2015). SNPs under directional selection (threshold: > 5.4 M.XtX value) are shown in red between dashed lines.

### Habitat suitability modelling for the HNWA and FNEA groups

The indication of local adaptation in the HNWA and FNEA groups implies specific environmental requirements influencing their growth. To elucidate the potential geographical distribution of these groups, we conducted habitat suitability analysis using historical climate data and future climate models for these two groups (Figures 6 and 7). Cross-validation yielded AUC estimates exceeding 0.970 for both groups, indicating the models’ robust discrimination capability. Analyses based on historical climate data unveiled that the potential distributions of both the FNEA group (Figure 6A) and the HNWA group (Figure 7A) are confined to relatively small, specific areas on the globe. The most relevant factors influencing the FNEA group were Annual Mean Temperature (variable 1), Mean Temperature of Coldest Quarter (variable 11), Temperature Seasonality (variable 4) and Mean Temperature of Driest Quarter (variable 9), while Isothermality (variable 3) and Temperature Seasonality (variable 4) were identified as the key determinants for the HNWA group (Supplementary figure 3). Pairwise comparison of D and I indices applied to habitat suitability distributions between FNEA and HNWA were 0.1 and 0.4, respectively, confirming their differential geographical distribution (Supplementary Table 9A and B). The potential geographical distribution of these two groups of maize was also modelled employing two future climate scenario models, CNRM-CM6-1 (Voldoire et al., 2019) and MRI-ESM2-0 (Yukimoto et al., 2019), for the period 2081-2100 and under four CO_2_ emission scenarios (SSP5-8.5, SSP3-8.7, SSP2-4.5, and SSP1-2.6) (Figures 6B and 7B). Pairwise comparison of D and I indices applied to habitat suitability distributions between historical climate and future climate models were on average 0.21 (D) and 0.48 (I) for FNEA and 0.19 (D) and 0.48 (I) for HNWA, indicating a shift in the geographical distribution of both groups in future climate conditions (Supplementary Tables 9C, D, E and F, respectively). D and I indices comparing future climate models within themselves were on average 0.87 (D) and 0.9 (I) for FNEA and 0.75 (D) and 0.84 (I) for HNWA, showing high similarity in the outcomes of the different models for each group (Supplementary Tables 10C, D, E and F, respectively). The results of our modelling suggest that suitable areas for the HNWA will significantly decrease, almost disappearing, while areas with favourable conditions for the FNEA will expand, albeit shifting towards more tropical latitudes.

**Figure 6.**
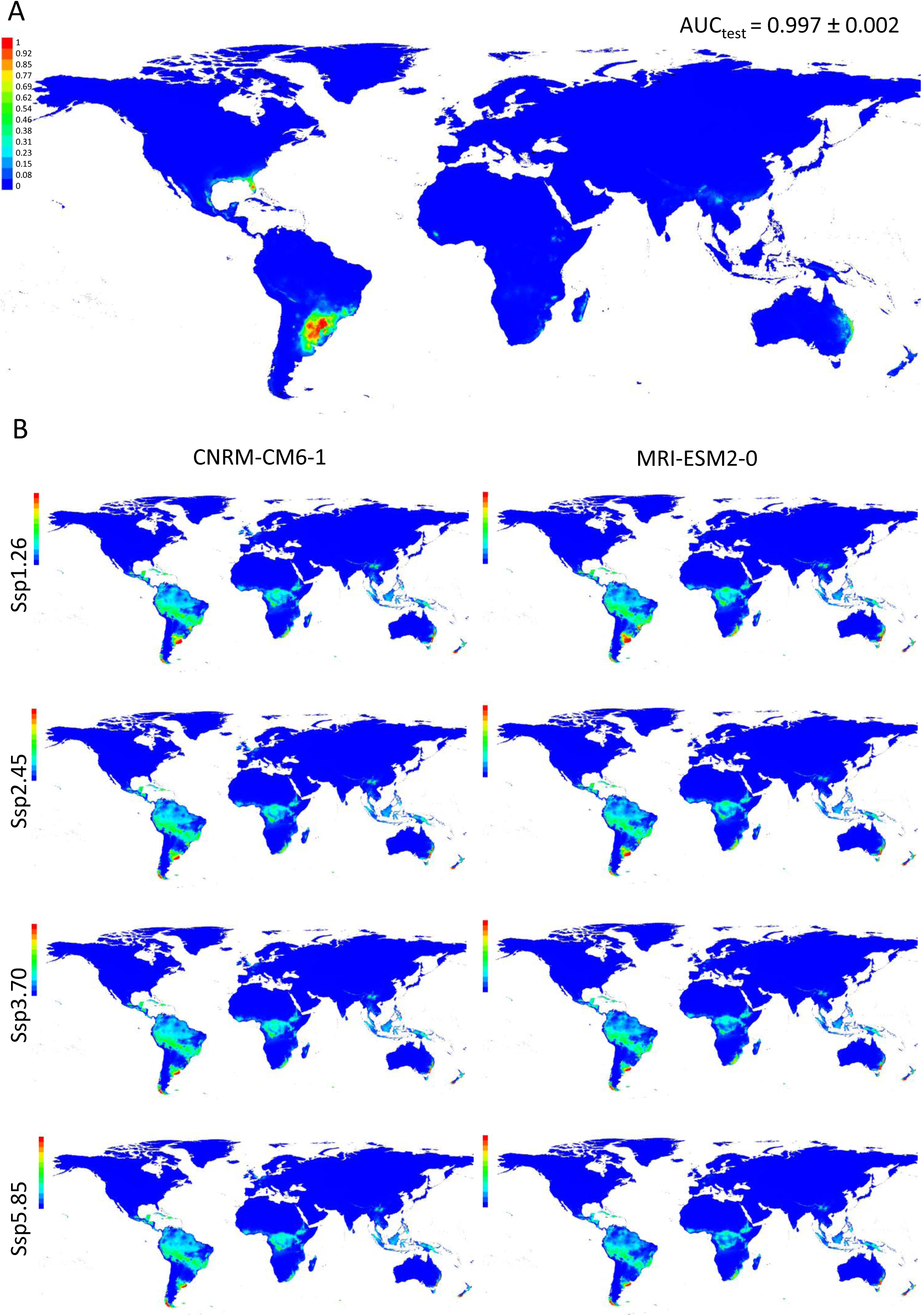
Habitat suitability modelling of Floury maize of Northeastern Argentina (FNEA) performed with MaxEnt (Phillips et al., 2004). Model in panel (A) represents the distribution of FNEA in the world employing altitude and historical climate data. This model was projected into two future climate scenario models, CNRM-CM6-1 (Voldoire et al., 2019) and MRI-ESM2-0 (Yukimoto et al., 2019), for the period 2081-2100 and under four CO_2_ emission scenarios (SSP1-2.6, SSP2-4.5, SSP3-8.7, SSP5-8.5) (B). The colours in the references indicate the strength of the prediction for each map pixel. The graphs show the average of 4 runs.

**Figure 7.**
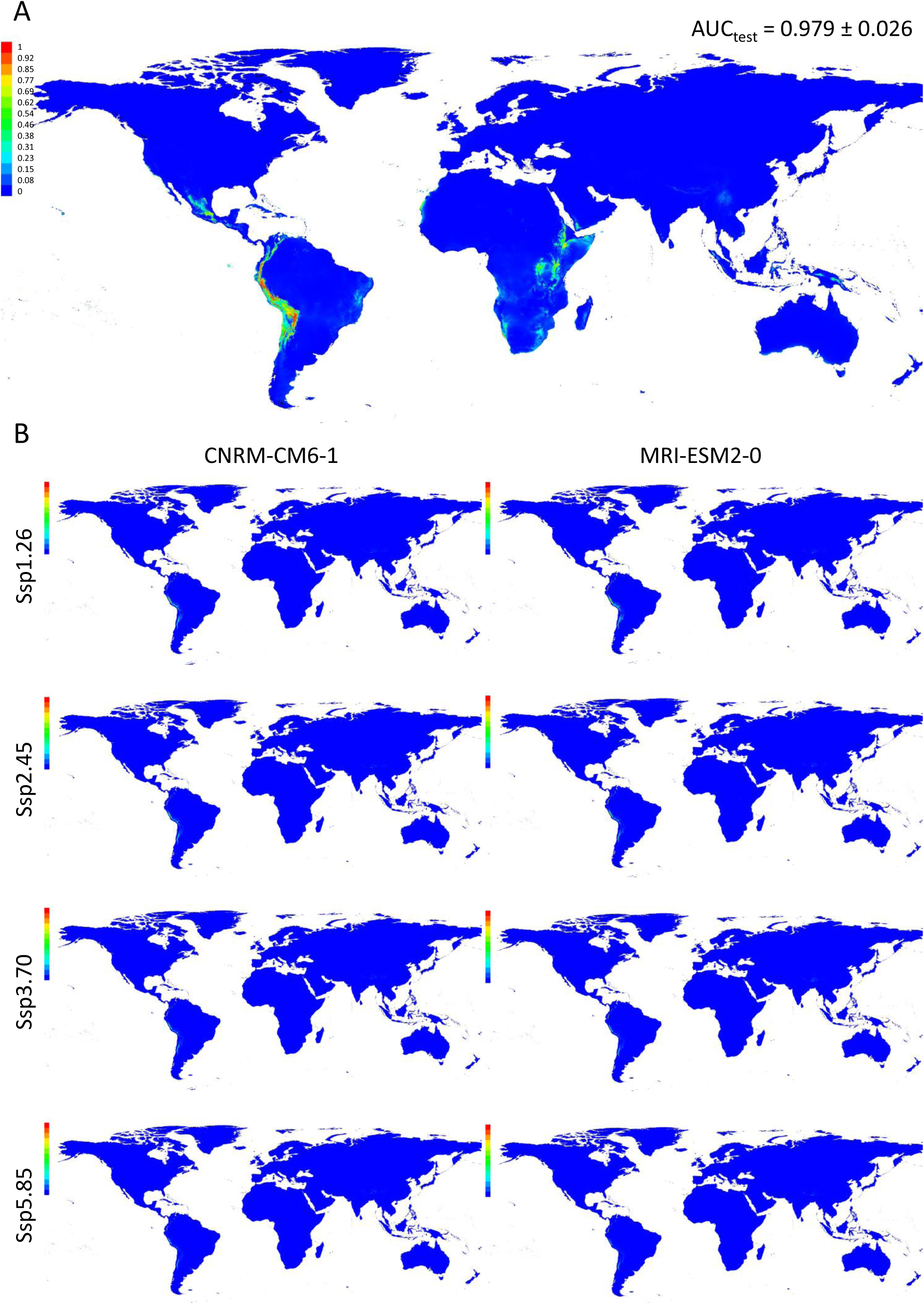
Habitat suitability modelling of Highland maize of Northweastern Argentina (HNWA) performed with MaxEnt (Phillips et al. 2004). Model in panel (A) represents the distribution of HNWA in the world employing altitude and historical climate data. This model was projected into two future climate scenario models, CNRM-CM6-1 (Voldoire et al. 2019) and MRI-ESM2-0 (Yukimoto et al. 2019), for the period 2081-2100 and under four CO_2_ emission scenarios (SSP1-2.6,SSP2-4.5,SSP3-8.7,SSP5-8.5) (B). The colours in the references indicate the strength of the prediction for each map pixel. The graphs show the average of 10 runs.

## Discussion

The delineation of evolutionary significant units is crucial for accurately interpreting EBVs. A priori delimitation of the groups examined in this work was based on genetic evidence derived from microsatellite markers and further supported by plastome sequences, morphological and phenological traits (Lia et al., 2009; Bracco et al., 2016; López et al., 2021; Rivas et al., 2022). However, by assessing genome-wide genetic diversity, we aimed at enhancing resolution, while simultaneously exploring both neutral and adaptive variation. Consistent with the findings of Rivas et al. (2022) and Bracco et al. (2016) concerning NWA, our SNP data demonstrate a clear separation among floury landraces cultivated above 2,000 m.a.s.l. (HNWA), floury landraces cultivated below 2,000 m.a.s.l. (LNWA), and popcorn landraces (PNWA) (Figure 2). While the HNWA group exhibited notable cohesion, individuals from LNWA and PNWA displayed relatively high levels of admixture and lacked well-defined clusters in the multivariate analyses. Moreover, our population structure results further confirmed the presence of two distinct groups in the Northeast of Argentina, FNEA and PNEA, as documented by Bracco et al. (2012, 2016), with the FNEA group consistently identified across various analyses (Figure 2).

As previously highlighted, Bracco et al. (2016) demonstrated the inclusion of HNWA maize within the Andean cluster defined by Vigouroux et al. (2008). Additionally, PNWA maize exhibited a close affiliation with landraces from Highland Mexico and Southern U.S. (Bracco et al., 2016). Conversely, FNEA and PNEA could not be linked to any other maize group within the Americas (Bracco et al., 2016). Likewise, the origins and affiliations of LNWA germplasm remain uncertain, and a direct comparison of this group with other lowland gene pools in South America had not been conducted prior to the present study. The degree of admixture inferred by STRUCTURE for LNWA, coupled with its overlap with individuals from other groups in clustering and ordination analyses (Figure 2), makes it challenging to establish the origin of this germplasm or determine whether it constitutes a single evolutionary unit. In the light of the most recent hypothesis on the diffusion of maize into South America (Vigouroux et al., 2008; Kistler et al., 2018), a plausible explanation for the observed pattern for LNWA is that it emerged as a consequence of secondary contact between Andean and lowland maize from eastern South America during pre-Columbian times. Alternatively, it could also be attributed to recent introgression between native landraces and improved germplasm derived from modern breeding. Indeed, the LNWA race *Orgullo Cuarentón* (Supplementary table 1), was classified by Cámara Hernández et al. (2012) as an incipient race with contributions from varieties developed in Argentina in the mid-1960s. It thus appears that further work in a global context is still needed to unveil the origin of LNWA.

In summary, guided by the outcomes of our population structure analyses, we concentrated on HNWA and FNEA to estimate EBVs and evaluate the conservation prospects of these two groups. Effective population size stands as a pivotal parameter in conservation genetics, as it governs the pace of allelic frequency changes due to genetic drift and informs on future levels of diversity (Hoban et al., 2022). Consequently, it is intricately associated with inbreeding and the depletion of genetic variation, in both neutral and adaptive loci (Allendorf et al., 2013). The contemporary Ne can be estimated using genetic data from a single sample (“population”) by calculating LD between loci (Waples and Do, 2010). Higher LD values signify smaller Nes, which could in turn imply that beneficial alleles are in linkage disequilibrium with deleterious ones, thereby potentially diminishing their positive effect on adaptation (Hoffmann et al., 2017). The observed extent of LD in FNEA and HNWA, 2.9 and 2.2 Mb, respectively (Figure 3), largely surpasses estimates previously reported for maize landraces (6.3 -30 Kb; Hufford et al., 2013; McLean-Rodriguez et al., 2021) and teosintes (*Zea mays mexicana*: 50 Kb, Z*ea mays parviglumis*: 10-22 Kb; Chen et al., 2022) but aligns closely with that of wheat landraces (3.6 Mb; Ma et al., 2022). In maize hybrids, LD blocks can average 28 Mb (Chaikam et al., 2019), while in rice hybrids, this figure can reach up to 75 Mb (Pradhan et al., 2020). The variations in the extent of LD between FNEA and HNWA result in a noticeable disparity in Ne, with estimated figures hovering around 50 individuals for FNEA and 200 individuals for HNWA (Figure 4C). Assessing the influence of methodological and/or biological factors, identified as potential distortions to Ne inferences based on LD, such as sampling, gene flow, or admixture (Gargiulo et al., 2023), poses challenges for our dataset. This complexity arises from the “populations” under scrutiny being somewhat abstract entities that represent diverse gene pools with dispersed geographical distributions. Nevertheless, although they should be taken with caution, these estimates offer a useful framework for interpreting the remaining EBVs and provide guidance for management actions. Consistent with a reduced Ne, individuals in the FNEA population demonstrate elevated inbreeding coefficients (F) (Figure 4B), rendering them more susceptible to inbreeding depression. This phenomenon, alongside its counterpart, heterosis, has proven to be notably significant in maize, as elevated F values have been associated with considerable yield reductions (Roff, 1997). Remarkably, genetic diversity estimates were found to be higher for FNEA compared to HNWA (Figure 4A), a result that might appear unexpected considering the differences in contemporary Ne. This discrepancy suggests that FNEA underwent a relatively recent bottleneck originating from an ancestral population that likely possessed greater diversity than HNWA. Changes in heterozygosity are not immediately evident following a reduction in population size (Keyghobadi et al., 2005; Lowe et al., 2005; Hoban et al., 2022). Conversely, the reduced variability observed in HNWA is consistent with the limited genetic diversity previously reported for the Andean group as a whole and is likely a consequence of the founder effect that led to the formation of this lineage (Vigouroux et al., 2008; Takuno et al., 2015; Bracco et al., 2016). It is noteworthy that both FNEA and HNWA, as well as the overall genome-wide diversity indices derived from this study, exhibit values at the lower spectrum of estimates reported for a diverse array of landraces and teosintes (Hufford et al., 2013; Rivera-Rodriguez et al., 2023), underscoring the vulnerability inherent in these groups. According to the estimates of Franklin (1980) and Soulé (1980) for natural populations of outbreeding species, a population should maintain a Ne of at least 50 individuals to avoid inbreeding depression in the short term. To minimise the impact of genetic drift and retain evolutionary potential, the Ne should surpass 500 individuals. Although specific Ne thresholds for cultivated plants remain undetermined, and annual species such as maize may tolerate lower Ne, the conjunction of high F and low Ne for FNEA suggests an elevated susceptibility to fitness and variability reductions (Hoffmann et al., 2017; Gaitán-Espitia and Hobday, 2021; Hoban et al., 2022). On the other hand, despite lower F values and higher Ne estimates for HNWA, this group may also encounter challenges in adapting to climate change, as indicated by low nucleotide diversity and Ne values below the recommended threshold of 500 individuals.

Divergence between populations, as measured by Fst indices, can account for the distinctiveness of each gene pool. The genome-wide Fst estimate for the HNWA-FNEA pair (Fst=0.07; Figure 5A) exceeded the values reported by Takuno et al. (2015) in their study comparing highland and lowland maize landraces from Meso- and South America (Fst=0.024 and 0.047). This higher Fst value suggests a more pronounced differentiation in allele frequencies between the highland and lowland germplasm of southern South America. This divergence can be attributed to smaller Ne, or more limited gene flow within the region.

It has been proposed that genetic variation of adaptive significance serves as a more reliable predictor of the long-term success of populations compared to overall genetic variation (Hoffmann et al., 2017; Kardos et al., 2021). To quantify adaptive differences, outlier detection methods come into play by identifying loci characterised by high genetic differentiation relative to the overall population structure, indicative of their likely involvement in divergent selection. The identification of selection signatures at multiple SNPs in the comparative analysis between HNWA and FNEA (Figure 5B), coupled with compelling evidence of local adaptation within Mexican and other South American maize landraces (Gates et al., 2019; McLean-Rodríguez et al., 2021; Wang et al., 2021; Janzen et al., 2022), suggests that these two groups exhibit signs of local adaptation.

Several studies have identified a correlation between flowering time or anthesis/silking interval and local adaptation in maize landraces (Mercer and Perales, 2019; Gates et al., 2019; Wang et al., 2021; Janzen et al., 2022; McLean-Rodriguez et al., 2021). The modification of flowering time through domestication has been crucial for extending the adaptability of various crops to diverse latitudes, a phenomenon also observed in wheat, barley, and rice (Nakamichi, 2015). In this study, two genes associated with flowering stand out among those containing outlier SNPs (Supplementary Table 8A). The first one, Zm00001d014690, known as *Arftf35* (ARF-transcription factor 35), encodes a protein involved in auxin-related axillary meristem formation in maize inflorescences (Galli et al., 2015; Galli et al., 2018). The second gene, Zm00001d015765, is an ortholog of Arabidopsis AtSWC4, which suppresses the expression of FT (florigen) and accelerates flowering time when knocked down (Gómez-Zambrano et al., 2018). Additionally, three outlier SNPs were found within genes whose expression is modified under stress conditions (Supplementary table 8A): the gene Zm00001d020497, identified as *cipk28* (calcineurin B-like-interacting protein kinase28), has been observed to exhibit responses to both salt and drought stresses (Chen et al., 2013; Feng et al., 2022). Similarly, Zm00001d047587 encodes a glucose-6-phosphate dehydrogenase (G6PDH3) and has demonstrated induction under osmotic and cold stress (Li et al., 2023). Furthermore, Zm00001d025651, orthologous to the *Arabidopsis* poly(A)-specific ribonuclease AtPARN, is implicated in a mRNA degradation system crucial to ABA, salicylic acid, and stress responses in *Arabidopsis* (Nishimura et al., 2005). These findings align with the concept that locally adapted landraces typically grow in marginal and stressful environments. Consequently, their adaptation may involve stress-related genes that contribute to fitness trade-offs (Corrado and Rao, 2017; VanWallendael et al., 2019).

Recent comparisons of genomic responses to selection have shown the participation of large haplotype blocks in population adaptation to new environmental conditions (Hoffmann et al., 2017). In this study, besides identifying outlier SNPs within genes, three chromosomal regions have emerged as potentially involved in local adaptation (Supplementary table 8B). The first spans positions 96,799,426 to 97,851,477 on chromosome 3. Structural variation analysis among the founders of the maize Nested Association Mapping (NAM) population revealed a large inversion encompassing this region, present in the inbred lines P39 and Oh43 (Hufford et al., 2021). Notably, this region has previously been associated with flowering time determination in both landraces (Navarro et al., 2017) and the NAM population (Buckler et al., 2009). Chromosomal inversions with adaptive significance may harbour genes influencing multiple traits (Huang and Rieseberg, 2020). Indeed, on chromosome 3, this region includes the *ys3* gene (Zm00001d041111, GRMZM2G063306), which has been shown to be under selection in *Z. mays* ssp. *parviglumis* (Aguirre-Liguori et al., 2017), and involved in iron homeostasis (Xu et al., 2022; Nozoye et al., 2013), a trait potentially important in the distinctive lateritic, iron-rich, red soils of NEA (Píccolo et al., 1998). Furthermore, the regions identified on chromosomes 7 and 10 (Supplementary table 8B) overlap with genomic tracts of *Z. mays* ssp. *mexicana* introgression into maize, previously associated with highland adaptation (Hufford et al., 2013, Calfee et al., 2021).

The distribution of genetic diversity is significantly influenced by geographic and climatic features, and the increasingly dynamic environmental conditions present a substantial threat to locally adapted germplasm. The potential distribution of the HNWA and FNEA groups under historical climatic conditions (Figures 6A and 7A) is in line with the limited distribution previously observed by Bracco et al. (2016). Utilising future climate scenarios in distribution models unveils potential risks to the persistence of these maize groups, particularly of HNWA (Figures 6B and 7B). As highland maize, HNWA faces greater environmental restrictions (Figure 7B), akin to predictions made for high-altitude teosintes (Ureta et al., 2012; Sanchez González et al., 2018; Aguirre-Liguori et al., 2019). The FNEA group, on the other hand, shows a projected displacement of suitable areas to other regions worldwide (Figure 6B). Range shifts due to climate change have been well-documented for numerous wild species (Wiens, 2016). For cultivated species like maize, the anticipated lack of suitable future climatic conditions in their regions of origin also poses a threat to the well-being of local communities. These findings underscore the importance of expanding research on how maize landraces will respond to climate change, incorporating not only local adaptation as a study variable but also considering plasticity.

## Conclusions

The genetic diversity of species allows them to adapt to environmental changes, evolve, avoid inbreeding depression, maintain fitness in their original environments and give rise to new species (Hoban et al., 2022). Assessing this diversity through various population genetics metrics, collectively termed EBVs by Hoban et al. (2022), provides insights into the status and trends of genetic variability. Our findings emphasise the necessity of treating FNEA and HNWA as distinct conservation units, highlighting an imminent risk of genetic diversity loss among maize landraces in northern Argentina. This concern is underscored by the low Ne values and elevated inbreeding coefficients observed in the FNEA group, coupled with low Ne values and diminished nucleotide diversity in the HNWA group. These indicators point towards ongoing or potential genetic erosion, constraining the adaptability of landraces to environmental variations. The swift pace of climate change poses an additional challenge, potentially hindering the evolution of these locally adapted landraces within their native environments (Aitken and Whitlock, 2013; Gaitán-Espitia and Hobday, 2021). Furthermore, species distribution modelling under future climate scenarios predicts a noticeable reduction in suitable cultivation areas. In conclusion, our results suggest that the long-term conservation of HNWA and FNEA landraces is jeopardised by the dual threats of genetic erosion and climate change. Therefore, it is imperative to promote their conservation both in situ and ex situ and expand the study of their plasticity and local adaptation to enhance our understanding of their environmental responses.

## Grant information

This research was funded by the Fondo para la Investigación Científica y Tecnológica (FONCYT, https://www.argentina.gob.ar/ciencia/agencia/fondo-para-la-investigacion-cientifica-y-tecnologica-foncyt), grants PICT 2013-0838, PICT 2016-1101, and PICT 2021-1286, and the Instituto Nacional de Tecnología Agropecuaria (INTA, https://www.argentina.gob.ar/inta), grant 2023-PD-L01-I085.

## Competing interests

The authors have declared that no competing interests exist.

## Supplementary tables

**Supplementary table 1.** Data of individuals sequenced by ddRADseq. A priori classification was based on Lia et al. (2009), Bracco et al. (2016), López et al. (2021) and Rivas et al. (2022). Individuals unequivocally assigned to the FNEA and HNWA genetic clusters by STRUCTURE and DAPC methods (membership coefficients or assignment probabilities > 0.75) (Figure 2C and D) are marked in orange and green, respectively. FNEA: Floury maize of Northeastern Argentina. PNEA: Popcorn of Northeastern Argentina. HNWA: Highland maize of Northwestern Argentina. LNWA: Lowland maize of Western Argentina. PNWA: Popcorn of Northwestern Argentina. VAV: ID of the “N.I. Vavilov” Plant Genetic Resource Laboratory, Faculty of Agronomy, University of Buenos Aires. ARZM: ID of the “Banco Activo de Germoplasma INTA Pergamino”. Coordinates are provided in decimal degrees.

**Supplementary table 2.** Unfiltered VCF file obtained with Stacks v1.42 (Catchen et al., 2013).

**Supplementary table 3.** Filtered VCF file. Filtering was performed with VCFtools (Danecek et al., 2011).

**Supplementary table 4.** Filtered and imputed VCF file. Imputation was carried out with Beagle (Browning et al., 2018).

**Supplementary table 5.** Filtered, imputed, and annotated VCF file. Annotation was performed with SnpEff (Cingolani et al., 2012).

**Supplementary table 6.** Occurrence locations of FNEA (Floury maize of Northeastern Argentina) and HNWA (Highland maize of Northwestern Argentina) individuals employed in the MaxEnt analyses. Locations were retrieved from Bracco et al. (2016) and this work (Supplementary Table 1). Groups were limited based on the STRUCTURE and DAPC analyses (membership coefficients or assignment probabilities > 0.75; Figure 2C and D). Duplicated occurrence locations were merged into one location. Coordinates are provided in decimal degrees.

**Supplementary table 7.** Estimated allele frequency (P) divergence among groups computed using point estimates of P by STRUCTURE. K=4. The classification of each group was based on the majority presence of groups defined a priori according to Lia et al. (2009), Bracco et al. (2016), López et al. (2021) and Rivas et al. (2022): FNEA (Floury maize of Northeastern Argentina); PNEA (Popcorn of Northeastern Argentina); HNWA (Highland maize of Northwestern Argentina); LNWA (Lowland maize of Western Argentina), and PNWA (Popcorn of Northwestern Argentina).

**Supplementary table 8.** (A) Supplementary data of the identification of outlier loci with BayPass (Gautier 2015), including SNP basic data, BayPass statistics information, SnpEff annotation of the found outlier loci, allelic frequencies of the outlier loci, and functional annotation of genes that contain SNPs identified as outlier loci in their bodies. HNWA: Highland maize of Northwestern Argentina. FNEA: Floury maize of Northeastern Argentina. (B) Identification of genes within 2 Mb intervals around outlier SNPs.

**Supplementary table 9.** Output of ENMTools (Warren et al., 2021) showing the Schoener’s D (Schoener et al., 1968) (A, C, E) and the I statistic (Warren et al., 2008) (B, D, F) comparing MaxEnt distributions for historical climate between FNEA (Floury maize Northeastern Argentina) and HNWA (Highland maize of Northwestern Argentina) (A, B) and between historical climate and future climate models for FNEA (C, D) and for HNWA (E, F). Green and purple indicate values that were averaged for comparison.

## Notes

### Competing Interest Statement

The authors have declared no competing interest.

