## Supplemental figures 1 to 3 for "Genome-wide diversity in lowland and highland maize landraces from southern South America: population genetics insights to assist conservation"

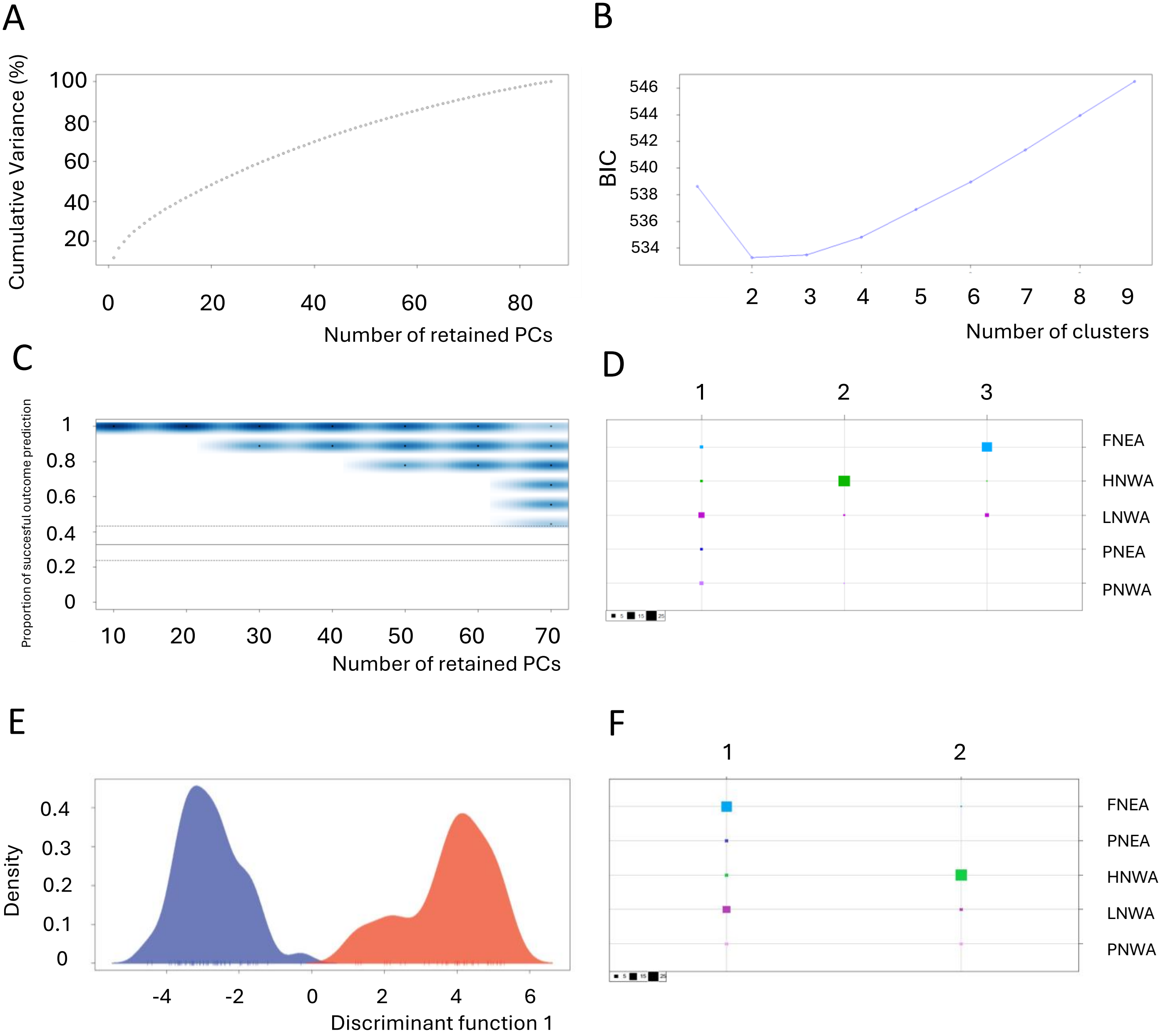


**Supplementary Figure 1.** Supplementary data of the Principal Component Discriminant Analysis (DAPC) performed with Adegenet in R (Jombart et al., 2008) and shown in Figure 2C. A) Variance explained by PCA (Principal Component Analysis). B) Values of BIC (Bayesian information criterion) versus number of clusters. C) DAPC cross validation. D) Contingency table of the K=3 DAPC (x-axis: DAPC groups, y-axis: maize classification, size of squares: number of individuals). E) Density graph for K=2. F) Contingency table of the K=2 DAPC (x-axis: DAPC groups, y-axis: maize classification, size of squares: number of individuals). HNWA: Highland maize of Northwestern Argentina. LNWA: Lowland maize of Western Argentina. PNWA: Popcorn of Northwestern Argentina. FNEA: Floury maize of Northeastern Argentina. PNEA: Popcorn of Northeastern Argentina. Total number of individuals: 87.

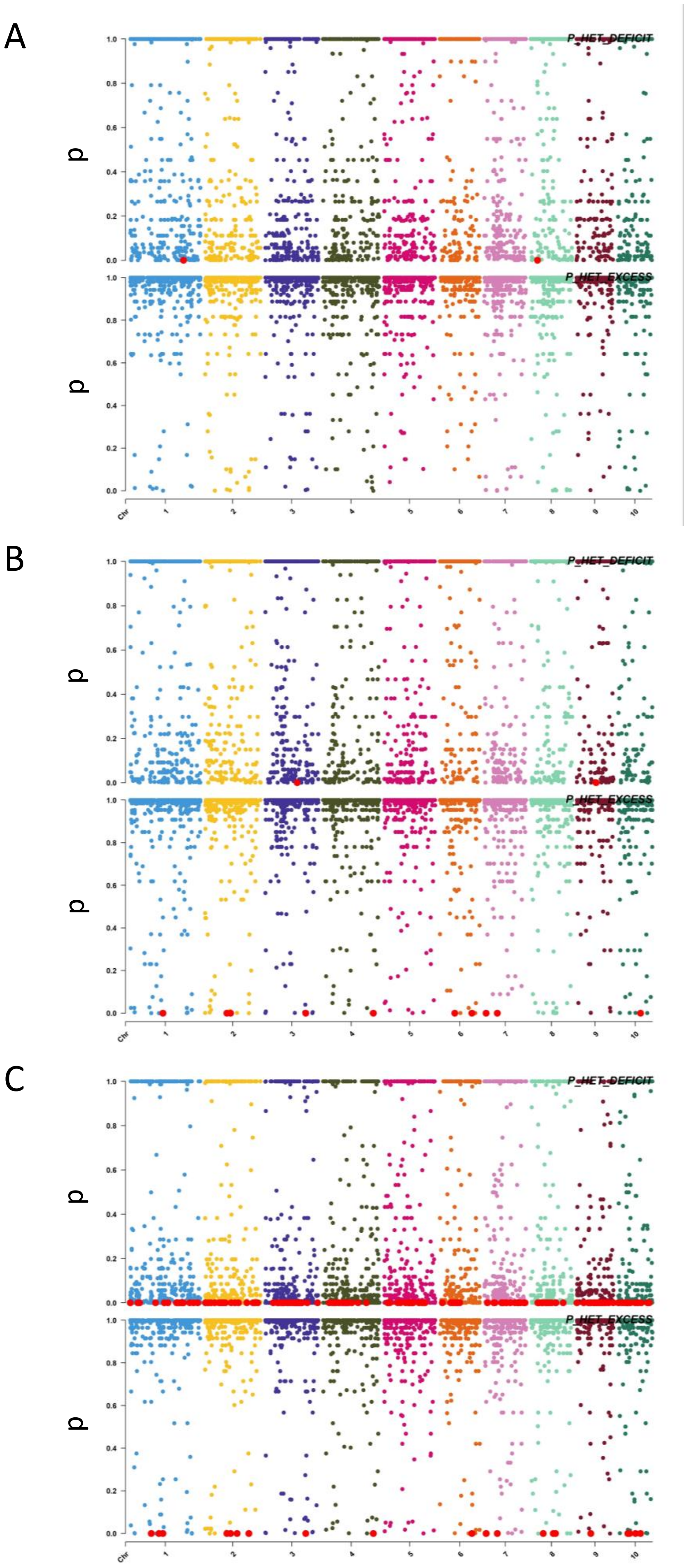


**Supplementary figure 2.** Hardy-Weinberg equilibrium obtained with VCFtools (Danecek et al., 2011) in (A) Floury maize of Northeastern Argentina (FNEA), (B) Highland maize of Northwestern Argentina (HNWA) and (C) all individuals employing the $\chi^2$ test. Upper panel: excess heterozygotes. Lower panel: heterozygotes in default. The plots show the p-values versus SNP genomic positions. Red dots indicate statistically significant excess or defect heterozygotes (p-value < 1.42e-5; p-values corrected for multiple testing by the Bonferroni test).

A

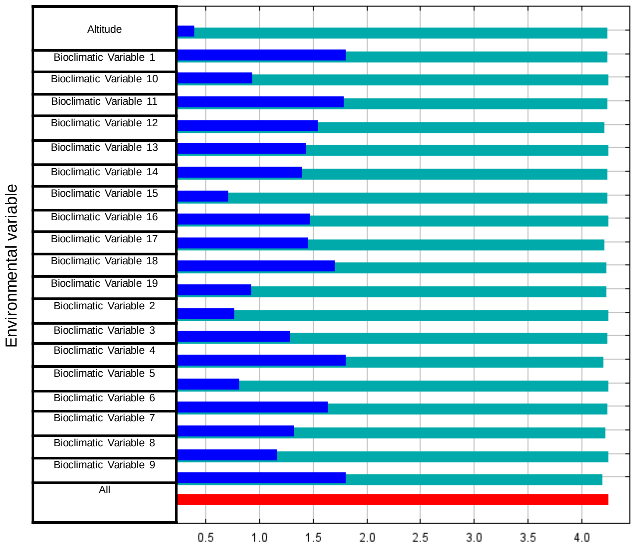


B

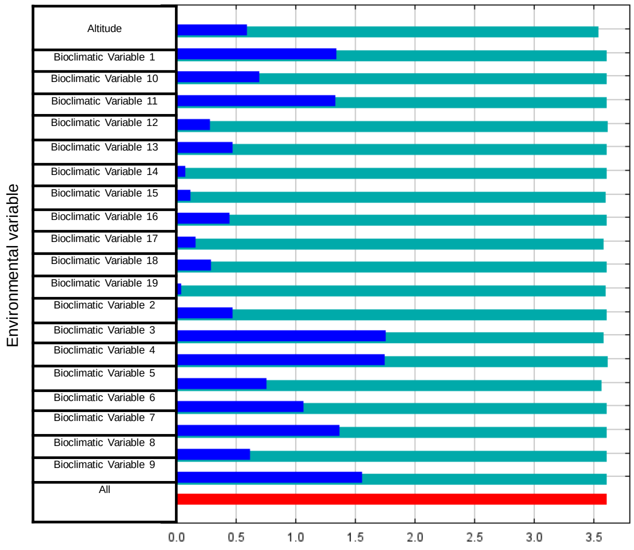


C

| Bioclimatic Variable | Definition |
|---|---|
| Bioclimatic Variable 1 | Annual Mean Temperature |
| Bioclimatic Variable 2 | Mean Diurnal Range (Mean of monthly (max temp - min temp)) |
| Bioclimatic Variable 3 | Isothermality (Bioclimatic variable 2/Bioclimatic variable 7) (×100) |
| Bioclimatic Variable 4 | Temperature Seasonality (standard deviation ×100) |
| Bioclimatic Variable 5 | Max Temperature of Warmest Month |
| Bioclimatic Variable 6 | Min Temperature of Coldest Month |
| Bioclimatic Variable 7 | Temperature Annual Range (Bioclimatic variable 5-Bioclimatic variable 6) |
| Bioclimatic Variable 8 | Mean Temperature of Wettest Quarter |
| Bioclimatic Variable 9 | Mean Temperature of Driest Quarter |
| Bioclimatic Variable 10 | Mean Temperature of Warmest Quarter |
| Bioclimatic Variable 11 | Mean Temperature of Coldest Quarter |
| Bioclimatic Variable 12 | Annual Precipitation |
| Bioclimatic Variable 13 | Precipitation of Wettest Month |
| Bioclimatic Variable 14 | Precipitation of Driest Month |
| Bioclimatic Variable 15 | Precipitation Seasonality (Coefficient of Variation) |
| Bioclimatic Variable 16 | Precipitation of Wettest Quarter |
| Bioclimatic Variable 17 | Precipitation of Driest Quarter |
| Bioclimatic Variable 18 | Precipitation of Warmest Quarter |
| Bioclimatic Variable 19 | Precipitation of Coldest Quarter |
| Altitude | Height above sea level |

**Supplementary figure 3.** Jacknife of regularised training gain for MaxEnt (Phillips et al., 2004) model of (A) Floury maize of Northeastern Argentina (FNEA) and for (B) Highland maize of Northwestern Argentina (HNWA) employing historical bioclimatic variables and altitudes from Worldclim (https://www.worldclim.org/data/cmip6/cmip6climate.html). Green: without variable. Blue: with only variable. Red: all variables. (C) Definition of the variables employed in the analyses.
